# A novel NO-dependent ‘molecular-memory-switch’ mediates presynaptic *expression* and postsynaptic *maintenance* of LTP in the octopus brain

**DOI:** 10.1101/491340

**Authors:** Ana Luiza Turchetti-Maia, Naama Stern-Mentch, Flavie Bidel, Nir Nesher, Tal Shomrat, Binyamin Hochner

## Abstract

The octopus brain shows a robust hippocampal-like activity-dependent LTP, which is NMDA-independent, yet associative and presynaptically expressed and, as shown here, also independent of protein synthesis. Have the molecular mechanisms for mediating this LTP evolved independently or have they converged? Here we report on a distinctive adaptation of the nitric-oxide (NO) system for mediation of the octopus LTP. Unlike the suggested role of NO in LTP *induction* in the hippocampus, in octopus, inhibitors of NO-synthase (NOS) did not block LTP *induction* but either 1) reversibly ‘erased’ LTP expression, suggesting that a constitutive elevation in NO mediates the presynaptic LTP *expression* or 2) ‘reversed’ LTP *induction* and *maintenance* because a second LTP could be induced after inhibitor washout. We therefore propose a protein synthesis-independent ‘molecular-switch’, whereby NO-dependent NOS reactivation maintains NOS in its active state. Thus, while the octopus LTP shows marked evolutionary convergence with LTP in vertebrates, an extreme molecular novelty has evolved to mediate it.

## Introduction

Octopuses and other coleoid (modern) cephalopods (squid, cuttlefish) provide an example of the independent evolution of complex cognitive behaviors in invertebrates^1,2^. The area in their central brain controlling their outstanding learning and memory abilities, the vertical lobe (VL)^3,4^, matches the generic connectivity scheme of fan-out fan-in association/classification neural networks^5,6^ (Fig. 1a). In the octopus, the first fan-out synaptic layer shows a robust hippocampal-like activity-dependent LTP^5,7^. This LTP was shown to be important for consolidation of short- and long-term associative memories^4^.

The LTP of the octopus VL is NMDA-independent and involves a robust presynaptic facilitation of transmitter release at the first glutamatergic synaptic layer^5,7^ (Fig. 1a). Despite being NMDA-independent, the VL LTP demonstrates the essential associative induction properties of *specificity*, *cooperativity* and *associativity*, suggesting a Hebbian induction mechanism^7^. This makes the octopus VL LTP an example of an NMDA-independent, presynaptically expressed LTP, which best resembles the LTP in the mammalian mossy fiber – CA3 neuron synapses^8,9^. The presynaptic LTP at the hippocampal mossy fiber terminals is a textbook example for a non-associative LTP^10^, and indeed there are no unequivocal indications for postsynaptic dependence^9^. In contrast, the presynaptic LTP in the octopus VL represents a classic case of Hebbian induction of an exclusively presynaptically expressed LTP^7^. The prevailing dogma is that such a Hebbian presynaptic LTP induction mechanism requires a ‘Hebbian detector’ in the postsynaptic cell and a retrograde signal that moves from the post-to the presynaptic terminal where it mediates LTP induction. Consequently, a biologically active and readily diffusible molecule like nitric oxide (NO) and the enzyme that produces it, nitric oxide synthase (NOS), are natural candidates for mediating this form of LTP^11,12^.

The role of NO as retrograde messenger in LTP has been somewhat controversial^13^ since its role as a retrograde message was first convincingly demonstrated in cultured hippocampal neurons^14,15^. Recent studies have provided good evidence supporting NO as a retrograde messenger in a specific form of the presynaptic component of the hippocampal CA3-CA1 LTP. As in the octopus, this presynaptic component of the LTP is NMDA-independent and is induced by a stronger postsynaptic response that leads to Ca^2+^-dependent activation of NOS, likely through a voltage-dependent opening of L-type Ca^2+^ channels^16^. The generated NO diffuses to the presynaptic terminals, where LTP is induced by the canonical NO-dependent activation of the cGMP second messenger cascade^16–20^.

Considering the octopus LTP as a case of convergent evolution and that NO is a well-known mediator of synaptic plasticity in mollusks^21–24^ prompted us to test whether NO is also involved in the LTP of *Octopus vulgaris.* Moreover, previous works have suggested that NO is required for visual and tactile learning in the octopus^25–27^. Indeed, we found that the NO system is involved in the octopus LTP. However, our results also show that the nitrergic neuromodulatory system has undergone a major adaptation to achieve a novel molecular mechanism for mediating associative presynaptic LTP. This involves an ingeniously simple ‘molecular-memory-switch’ by which a single molecule, NO, mediates both the presynaptic *expression* of LTP and the very long-term *maintenance* through NO-dependent reactivation of NOS in the postsynaptic cell. This dynamic molecular-switch maintains LTP for a long time (>10 h), even when protein synthesis is blocked.

## Results

### The neuropils of the lobes associated with learning and memory stained intensely for NOS activity

We first tested the VL for NOS activity histochemically using the NADPH-diaphorase method^28,29^. There was a similar pattern of intense labeling in the VL and the subfrontal lobe (SuFL), areas associated with visual and tactile learning, respectively, which share a similar anatomical organization^30^ (Fig. 1e,f). Dense labeling was found in the inner neuropil zones (Fig. 1f,g), which contain the synaptic connections between the amacrine cells (AM) neurite terminals and the large efferent neurons (LNs) dendrites. The outer neuropil was more sparsely stained because in this region unstained axons that run in the SFL tract make sparse *en passant* connections with AM neurites. The AM neurites can be seen crossing the tract in faintly stained AM trunks (arrows; Fig. 1f), suggesting the presence of NOS in the AMs. The tiny AM cell bodies are not stained, likely because they are fully occupied by their nuclei^31^ (Stern-Mentch N. thesis).

### NOS inhibitors blocked LTP *expression* but not LTP *induction*

VL slice preparations were used to test whether NOS inhibitors affect LTP induction (Fig. 1b). As previously described^7^ and similarly to hippocampal slice preparations, we placed the stimulating and recording electrodes on the SFL tract at a short distance from each other. The tract was stimulated usually with paired test pulses and the evoked presynaptic tract potential (TP; Fig. 1b inset) and the postsynaptic field potential (fPSP) were recorded. The slices were exposed to NOS inhibitors for 30 minutes before LTP induction by high frequency stimulation (HF, four trains of 20 pulses at 50 Hz, 10 s inter-train interval). Fig. 1b shows that HF stimulation in the presence of L-NNA (10 mM) induced only small facilitation. However, following drug washout, the fPSP amplitude recovered to a much higher level than the control, and a second HF stimulation did not lead to further facilitation, indicating that LTP induction was fully activated in the presence of the blocker. These results led us to postulate that NO may be involved in LTP *expression* rather than LTP *induction*.

To test this possibility we added NOS inhibitors after LTP induction. While L-NAME (10 mM) completely inhibited the facilitated fPSP (Fig. 1c), this effect was absent when introducing the inactive enantiomer D-NAME (10 mM) as a control for specificity of NOS inhibition (Fig. 1d). A second HF stimulation before rinsing out the drug did not overcome the inhibition (Fig. 1c). The third HF, during recovery from the inhibition, showed no residual LTP, again demonstrating recovery of *expression* rather than inhibition of LTP *maintenance*. Both L-NAME and D-NAME had a similar effect on the amplitude of the TP (Fig. 1c,d insets), in contrast to L-NNA (Fig. 1b inset), indicating that the effects were not mediated by NO.

**Fig. 1.**
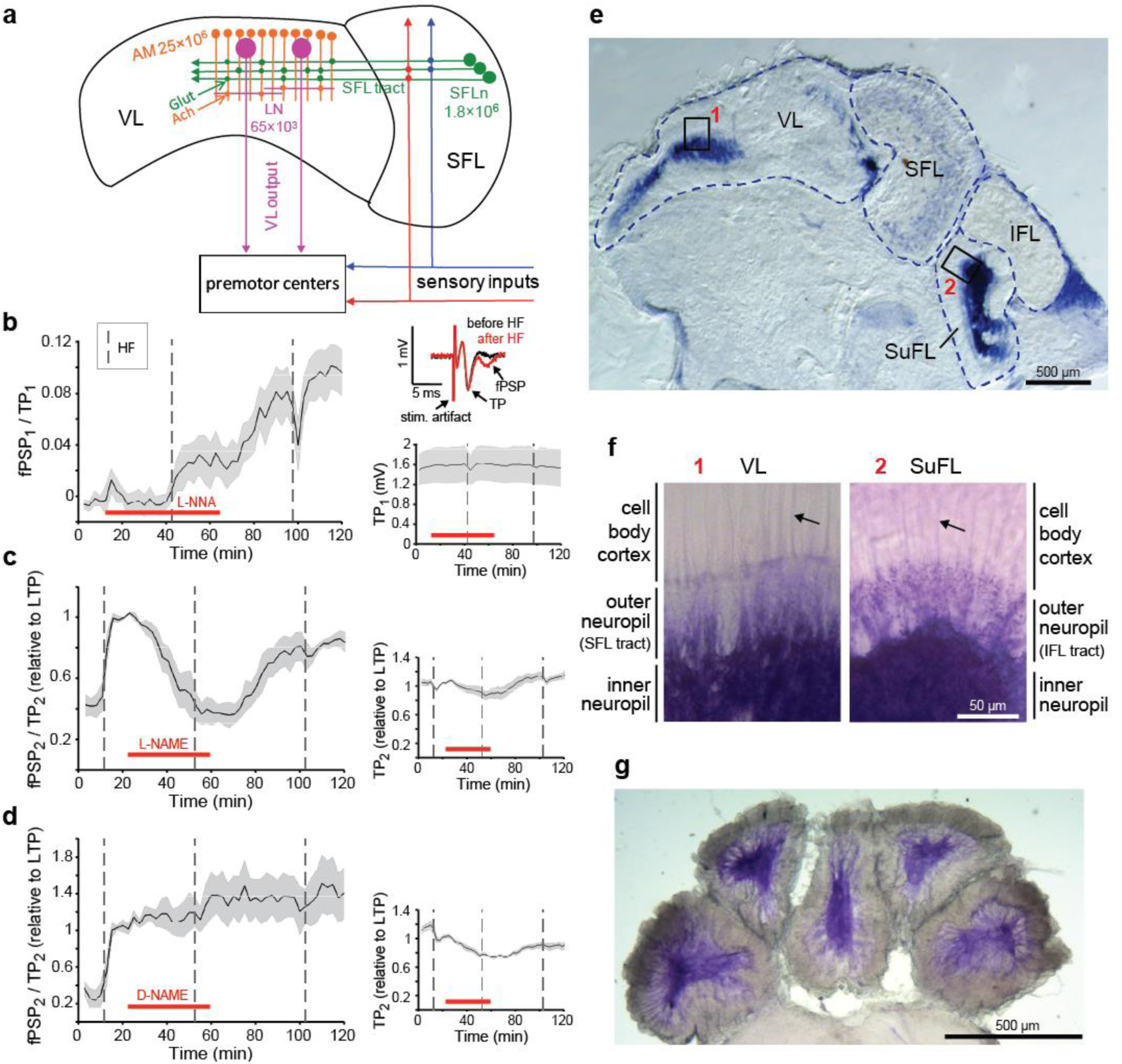
NOS inhibitors block LTP expression but not induction. **a** Connectivity scheme of the SFL-VL network. **b-d** The effects of NOS inhibitors (**b**, L-NNA 10 mM, *n*=7; **c**, L-NAME 10 mM, *n*=4) and **d**, an inactive enantiomer (D-NAME 10 mM, *n*=6) on the fPSP amplitude recorded in VL slice preparations. Data shown as mean ± s.e.m., dashed vertical lines mark HF stimulation, red bars mark drug exposure time. **e-g** Histochemical NADPH diaphorase staining of slices of the supraesophageal brain mass. **e** Sagittal section; note intense labelling of the neuropil in VL (1) and SubFL (2). **f** Higher magnification of the respective areas in **e**. Arrows point to labelling of AM trunks. **g** Transverse section of the five lobuli composing the VL. VL = vertical lobe; SFL= superior frontal lobe; SubFL = subfrontal lobe; IFL = inferior frontal lobe.

### NOS inhibitors block the presynaptic expression of LTP

The octopus VL LTP is mediated mostly, if not entirely, by an exceptionally large increase in the probability of transmitter release^7^, a process that is clearly manifested in large changes in the paired-pulse facilitation (PPF). In PPF, the first stimulus of a pair facilitates the synaptic release evoked by the second stimulus; the phenomenon is thought to be mediated by the residual Ca^2+^ entering the presynaptic terminals during the first stimulus^32^. We therefore employed a PPF analysis of the experiments shown in Fig. 1c to evaluate the mechanisms of inhibition.

The example in Fig. 2a,b shows the changes in the amplitudes of the first (fPSP1) and the second (fPSP2) fPSPs together with the PPF index (i.e., (fPSP2/TP2-fPSP1/TP1)/fPSP2/TP2) — note that this index equals 1 when fPSP1=0, as frequently happens in the controls). In the control, fPSP1 is much smaller than fPSP2 (A in Fig. 2a,b), and the PPF index is therefore high (0.85), indicating a large short-term facilitation (∼9-fold) of fPSP2 relative to fPSP1. Following LTP induction (B in Fig. 2a,b), both fPSPs are highly facilitated, but the relative increase of fPSP1 is much larger (∼21-fold) than that of fPSP2 (∼5-fold). This difference in the relative increase of the fPSPs is expressed in a sharp drop of the PPF index (B in Fig. 2a,b). Such a drop would not be expected were LTP mediated, for example, by an increase in postsynaptic responsiveness^33^, thus confirming the presynaptic origin of the VL LTP^7^.

The potentiation of both fPSPs was completely ‘erased’ following exposure to L-NAME (C in Fig. 2a,b), and the PPF index returned to control levels, indicating a reversal of the probability of release to the pre-LTP level. Following recovery from inhibition, the PPF index increased towards that of LTP levels (D in Fig. 2a,b), showing a reversible block of presynaptic facilitation. Fig. 2c shows similar behavior averaged from four such experiments. In conclusion, these reversible changes in PPF strongly suggest that NOS inhibitors specifically block LTP expression and that the elevation in NO concentration mediates the increase in probability of release from the SFL axonal terminals.

Is NO similarly involved in the late phases of LTP? We exposed the slices to an NOS inhibitor at different intervals after LTP induction (Fig. 2d-f). L-NNA (10 mM) could block LTP expression for at least 4 h after LTP induction. Similar to Fig. 2a-c, and irrespective of time after LTP induction, the PPF index was correlated with the level of fPSP inhibition. This result indicates that constitutive elevation of NO concentration is likely the main mechanism mediating the very long-term expression of the VL LTP.

**Fig. 2.**
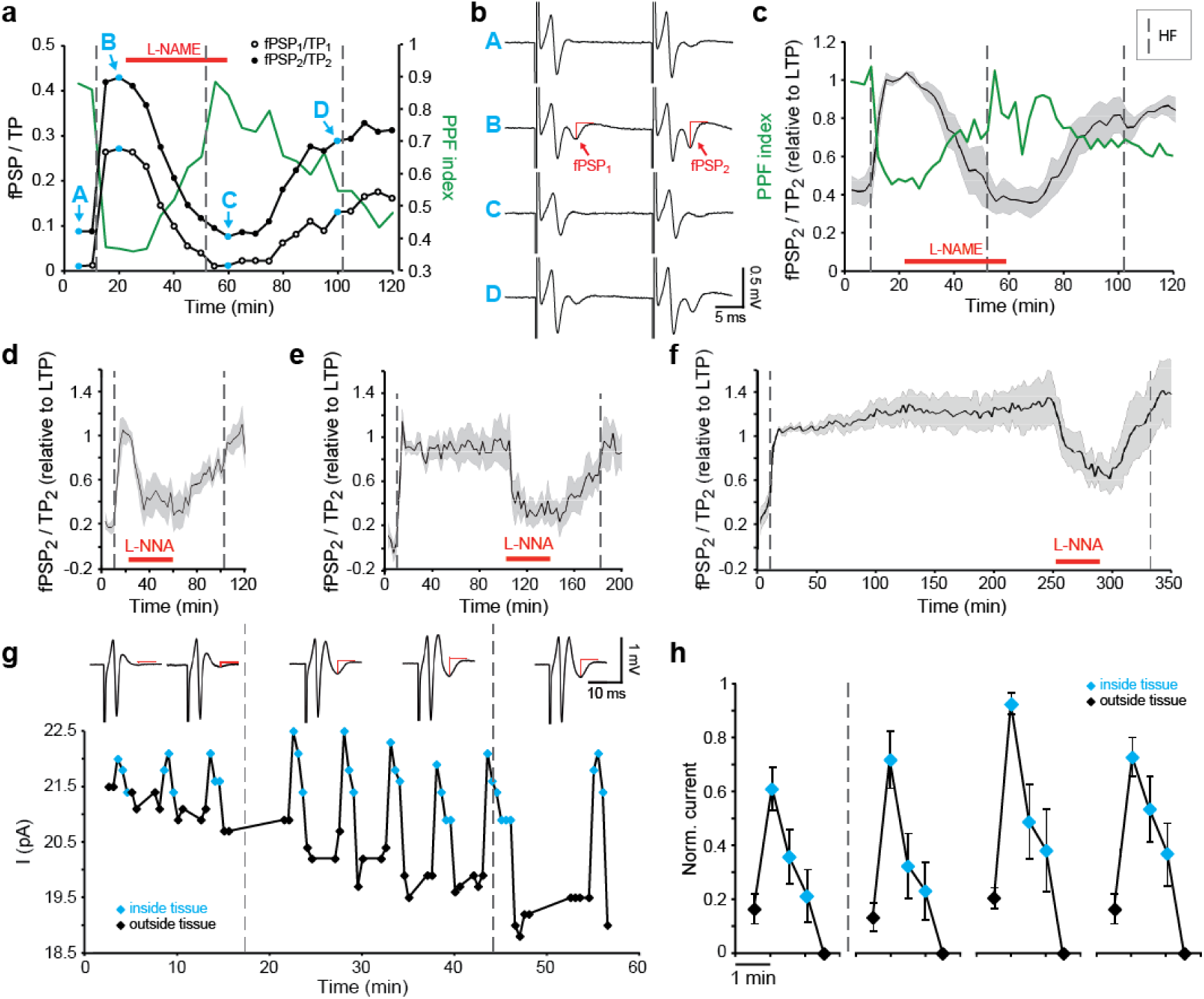
Constitutive elevation in NO concentration mediates the presynaptic expression of LTP. **a** Single experiment exemplifying the changes in the first (open circle) and second (filled circle) fPSP shown together with PPF index (green line). **b** Average traces at specific recording times in **a**, marked by corresponding letters (A, B, C, D). **c** Average of fPSP_2_/TP_2_ (*n*=4) shown together with average PPF index (green line) suggesting presynaptic origin of both LTP expression and NOS inhibition. **d-f** L-NNA (10 mM) blocks LTP irrespective of time after LTP induction: **d**, 10 min (*n*=4), **e**, 90 min (*n*=5), and **f**, 240 min (*n*=3). **g-h** Amperometric NO measurements in the VL of isolated brain preparations. **g** Amperometic signals during repeated insertions of the CFM into the VL neuropil. The corresponding fPSPs are shown in upper insets (red lines mark fPSP amplitude). The second HF stimulation was delivered when the electrode was inside the tissue. **h** Population response before and after HF stimulation (*n*=5). Data shown as mean ± s.e.m., dashed vertical lines mark HF stimulation, red bars mark drug exposure time.

### LTP maintenance involves constitutive elevation in NO concentration in the VL neuropil

To directly test whether maintenance is mediated by long-term elevation in NO concentration we adopted an electrochemical approach for measuring the extracellular concentration of NO amperometrically^34,35^ (Fig. 2g-h). A specially prepared NO-sensitive carbon fiber microelectrode (CFM, see Methods) was inserted for short periods into the VL neuropil. Following LTP induction, the clamping current of the CFM at the NO oxidation potential (750 mV) increased significantly and in parallel to the increase in fPSP amplitude (Fig. 2g, inset traces) to about 0.5 µM (the calibration of the electrode after the experiments in Fig. 2g, for example, indicated a sensitivity of about 6 pA / µM NO), which is a rather high biological NO concentration^36^. Typical for LTP saturation, both fPSP and NO signals were not further increased by a second HF (Fig. 2g).

Figure. 2h shows the average of five such experiments on isolated brains. NO concentration increased significantly after HF stimulation, with indication of a delayed elevation in NO concentration in the second measurement after LTP induction. Although we could show that, as expected, NOS inhibitors blocked the amperometric response together with blocking LTP expression, we found that NOS inhibitor also blocked the CFM response to NO (released from NO donor SNAP), and therefore we could not confirm that NOS inhibition effects are mediated via reduction in NO concentration.

### NO-dependent cGMP-cascade is likely not involved in VL LTP

A battery of pharmacological tools was used to test whether NO increases the presynaptic probability of transmitter release via the common cGMP pathway. We first tried to block the NO-dependent activation of soluble guanylyl cyclase (sGC). Neither ODQ (50 μM) nor methylene blue (0.1 mM) affected the LTP. Attempts to activate the downstream cGMP-dependent protein kinase with cGMP analogues dBcGMP (0.2 mM), 8-Br-cGMP (0.1 mM) and 8-pCPT-cGMP (10-100 μM) also did not facilitate the fPSP. Furthermore, the cyclic nucleotide phosphodiesterase inhibitor IBMX (1 mM) and the cGMP specific phosphodiesterase inhibitor ZAP (0.1 mM) had no facilitatory effect on LTP induced by HF or by a moderate rate of stimulation that induced a slow development of LTP. On the contrary, we have already shown that IBMX greatly suppressed transition to the long-term maintenance phase of LTP^37^. Taken together these experiments do not support the involvement of the cGMP cascade in the octopus VL LTP.

Drugs that modulate NO concentration levels, like the NO donors SNAP (1-100 μM), DETA (0.4 mM) and DEA (0.1 mM), also did not facilitate transmitter release, and the NO scavenger PTIO (1 mM) did not reduce release probability of control or of potentiated fPSP. If NO indeed participates in the VL LTP, then one explanation for these negative findings is that NO functions at a relatively high concentration as indicated by the amperometric results (see Discussion).

### The VL LTP is independent of *de novo* protein synthesis

Numerous studies indicate that the universal mechanism for maintaining the long-term phases of LTP involves *de novo* protein synthesis^38^. Surprisingly, here, neither LTP induction nor its very long maintenance appears to involve protein synthesis. The administration of 20 µM anisomycin for 2.5 h, starting 30 min before the induction of LTP, blocked neither induction nor maintenance of LTP for at least 10 h after LTP induction (Fig. 3a). Importantly, in four experiments exposing *in vitro* slices to 20 µM anisomycin dramatically suppressed protein synthesis, including the intense protein synthesis in the VL (Fig. 3c).

**Fig. 3.**
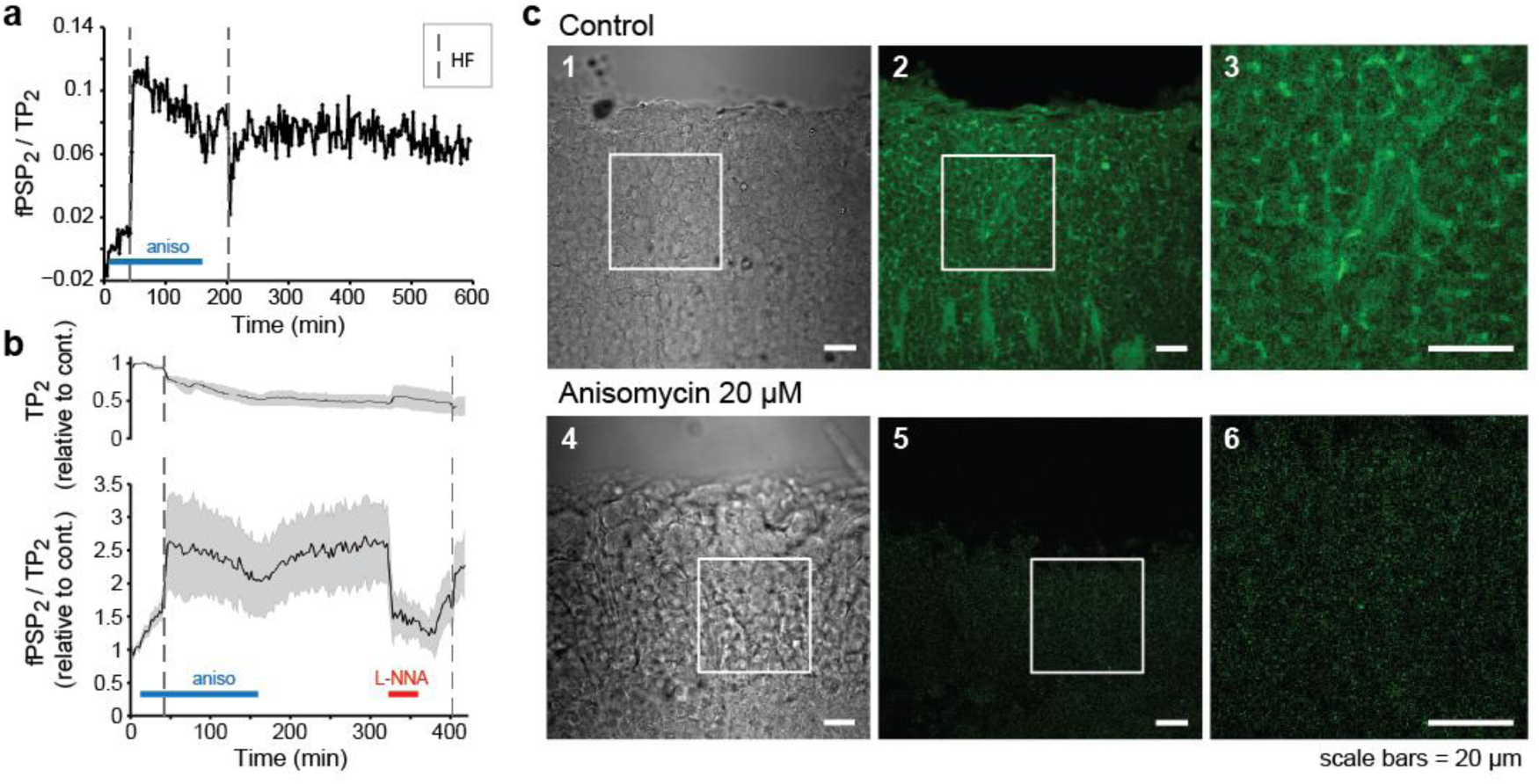
The VL LTP is protein synthesis-independent. **a-b** Testing the effect of exposure to anisomycin (20 µM, blue bars) on VL LTP induction and maintenance. **a** Example of a very long duration experiment (10 h). **b** Population average of four experiments (mean ± s.e.m.) where L-NNA (10 mM, red bar) blocked the anisomycin-resistant LTP (lower graph), and anisomycin affected the TP (upper graph). Dashed vertical lines mark HF stimulation. **c** Confocal microscopy images of *de novo* protein synthesis using the FUNCAT assay: 1,4 transmission bright-field images of the AM area; 2-3,5-6 fluorescent signal, largely suppressed by anisomycin. The white squares in 1-2 and 4-5 mark the areas of magnification displayed in 3 and 6, respectively.

### Long-term LTP maintenance involves NO-dependent NOS reactivation

The analysis of the kinetics of LTP inhibition provided an insight into the mechanisms of LTP maintenance. Fig. 4a,b shows two examples of the time course of L-NNA effects. The blue curve in Fig. 4a shows a gradual and only partial blockage of the LTP followed by a slow recovery during washout. An HF stimulation 40 min after onset of drug washout did not affect the slow recovery rate. In contrast, as shown in red, LTP expression was completely blocked within a few minutes of onset of the L-NNA effect, followed by small recovery during washout. Here, HF stimulation re-induced a robust second LTP that reached the pre-blocked level, suggesting that in this mode of inhibition both LTP maintenance and induction were ‘reversed’ to a pre-LTP level. The time courses of the population response of blocking and recovery (Fig. 4b) can clearly be segregated into two classes: one showing a simple reversible inhibition of LTP expression, the other presenting an irreversible inhibition of both expression and maintenance and reversal of induction.

**Fig. 4.**
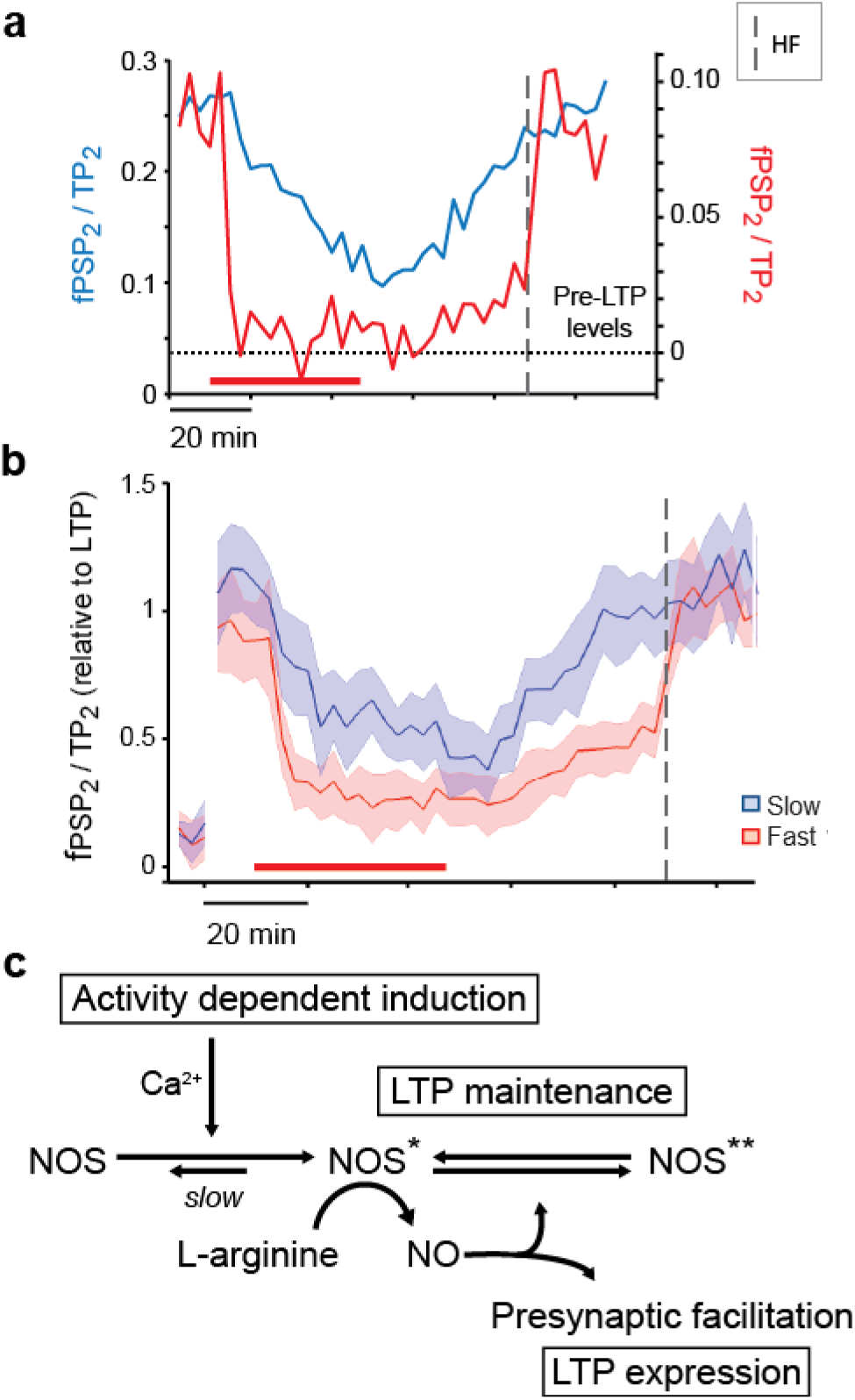
NO mediates both LTP expression and maintenance. **a** Examples of a slow and reversible L-NNA blockage of LTP expression (blue line) and of a fast and irreversible blockage of LTP expression and maintenance (red line). Note that only in the latter case did a second HF reintroduce LTP. **b** Population response classified according to these two groups (slow *n*=6, fast *n*=6). This data set was extracted from the recordings in Fig. 2d-f. **c** Proposed model for a dynamic molecular-switch mechanism, whereby NO mediates the increase in probability of presynaptic transmitter release and at the same time “locks” NOS in active states. (The Ca^2+^ dependence is based on Hochner et al.^7^).

## Discussion

Our results suggest the independent evolution of a novel and simple mechanism mediating an associative (Hebbian) LTP that involves postsynaptic induction and exclusively presynaptic expression – a form of LTP that has been highly anticipated following the discovery of NO as retrograde messenger. Presynaptic LTP is considered to provide a larger dynamical range of modulation relative to the postsynaptically expressed LTP (commonly mediated via NMDA dependent mechanism)^19^. This property is particularly characteristic of the octopus VL, where the probability of release (especially of fPSP1) is very low and it undergoes a huge presynaptic facilitation after LTP induction (e.g., 21 fold; Fig. 2a,b). That is, our results suggest that in the octopus VL an efficient and powerful molecular mechanism has evolved to enable this unique form of LTP. This was achieved by adapting the NO system to mediate LTP *expression* and *maintenance* while in the currently known NO-dependent presynaptic LTP, NO is involved in LTP *induction* via the canonical NO dependent cGMP second messenger cascade^11,12,39^.

Taken together our results lead us to suggest a simple molecular-switch mechanism that comprehensively explains all the properties of the VL LTP and is summarized schematically in Fig. 4c. The NOS inhibitor (L-NNA) reversibly blocks the conversion of L-arginine into NO, thus reducing LTP expression. In the fast and irreversible inhibition mode, a regenerative synergy between reduction in NO concentration and the concomitant reduction in NO-dependent NOS-reactivation leads to a faster, complete and irreversible deactivation of NOS itself. The return of NOS to its inactivated state allows a new induction of LTP by HF stimulation. The spontaneous recovery of NOS activity following the drug washout is due to a residual unblocked NOS-activity which is enough for a regenerative, NO-dependent, full NOS reactivation. Note that the positive-feedback properties of this molecular cascade explain inhibition of *expression* alone or blocking both *expression* and *maintenance*. Yet, it is also possible that each inhibition mode is associated with different types of synapses as we found, for example, with respect to the dependence of LTP induction on the postsynaptic response^7^.

Although the exact biochemical mechanisms involved are still not entirely clear, our findings are reminiscent of the molecular-switches based on kinase autophosphorylation suggested for hippocampal LTP^40,41^. Still, our findings suggest a simpler dynamical molecular-switch that can flip rapidly between ON and OFF states due to a positive-feedback loop (resembling the kinetics of non-inactivated voltage-gated ion channels). As originally suggested^40^, such a molecular-switch, which is based on covalent modifications of existing molecules, can provide the means for consolidation of long-term memory irrespective of *de novo* protein synthesis, as indeed we found in the octopus VL. The VL LTP is important for long-term memory consolidation that occurs outside the VL^4^, and protein synthesis is important for long-term memory acquisition^42,43^. Consequently, the molecular memory switch in the octopus VL may persist only until completion of memory storage outside the VL. Because LTP expression and maintenance can in principle be switched OFF and ON again very rapidly, the molecular-switch may provide a convenient checkpoint for supervisory inputs to, for example, ‘veto’ LTP maintenance. Both the ability of IBMX to suppress transition to the long-term maintenance phase^37^ and similar preliminary results with octopamine^44^ support this idea.

For this molecular-switch to function in the framework of Hebbian induction it is reasonable to assume that NO does not activate NOS directly (or at least only at a very high concentrations), because otherwise LTP would also spread to neighboring synapses, obliterating the Hebbian property of *synaptic specificity*. This assumption is supported by finding that release of even relatively high NO concentrations from an NO donor (SNAP) did not induce LTP. Accordingly, and as previous results suggest^7^, LTP *induction* involves activity/Ca^2+^-dependent activation of NOS and possibly only the active form (NOS*) can be reactivated by NO (Fig. 4c). Such a molecular constellation will, on the one hand, lead to Ca^2+^-dependent NOS activation in the AMs, contributing to the postsynaptic component of a Hebbian induction mechanism. On the other hand, it ensures synaptic *specificity,* as only the presynaptic terminals in the immediate vicinity of the activated NOS in the postsynaptic cell will be facilitated retrogradely by NO^11,12^.

It still remains to determine the downstream biochemical cascade through which NO mediates its effects on transmitter release and on NOS reactivation. Currently, there are two main biochemical cascades known to mediate NO effects. The best documented involves NO-dependent activation of soluble guanylyl cyclase (sGC) in the cGMP/PKG cascade that functions at a very low NO concentration ranging from 100 pM up to 5 nM^36,39^. The second involves NO-induced covalent protein modification through a direct non-enzymatic s-nitrosylation of cysteine residues that requires a much higher NO concentration than the cGMP cascade^45,46^. We have found no indication for the involvement of the cGMP cascade in VL LTP. Moreover, NO donors and NO scavengers neither facilitated nor inhibited the fPSP, respectively. This suggests that these manipulations were ineffective in interfering with the physiological range of NO concentration in the VL, especially at the synaptic connections, where the effective concentration is likely the highest^12^. Thus, considering these negative results together with the amperometric measurements that indicate a µM range following LTP induction and during maintenance (Fig. 2g,h), the relatively high concentration of competitive NOS inhibitors (10 mM) and the intense NADPH-d activity (Fig. 1e-g), we raise the possibility that LTP expression in the VL is mediated by a non-enzymatic process and suggest s-nitrosylation as a possible candidate^46^. Auto s-nitrosylation of NOS has been reported, but in that case it inhibited NOS activity^47^. Conceivably, the higher concentration required for s-nitrosylation may suit better the locality of the retrograde message effect, ensuring synaptic *specificity*.

In conclusion, our results provide new physiological insights into the growing understanding that cephalopod evolution involved an outstanding flexibility in the selection of neuronal and morphological novelties. This evolutionary flexibility, at the neurophysiological level, has been demonstrated by the surprising finding of dichotomic differences in synaptic plasticity processes in the VLs of octopus and its phylogenetically close relative, the cuttlefish *Sepia officinalis*^5^. Especially interesting are preliminary results suggesting that NO and NOS activity are absent in the cuttlefish^48^. Recent genomic studies suggest that coleoid cephalopods are unique in adopting multiple regulatory mechanisms that allow more ‘modular’ developmental frameworks^49^. In addition, this group has a huge expansion of RNA re-editing sites that allow a unique post-translational modification of core gene products^50^. Such modular mechanisms may have also facilitated the selection of an alternative molecular mechanism for the mediation of LTP with specific properties suitable for achieving species-specific behavioral requirements.

## Acknowledgments

We thank Avraham Susswein and Yosef Yarom, for their comments on an earlier version of the manuscript, and Jenny Kien for editorial assistance. We thank Hadas Erez and Micha Spira for helping with the confocal images. A.L.T.M. received postdoctoral fellowships from Lady Davis and Golda Meir Foundations, and The Edmond and Lily Safra Center for Brain Science (ELSC). This research was supported by the Israel Science Foundation (1928/15 and 1425/11) and the US-Israel Binational Science Foundation (2011466), and was aided by the Smith Family Laboratory at HUJI.

## Author contributions

A.L.T.M., T.S. and B.H. designed the study; A.L.T.M. performed and analysed the physiological experiments with contribution of T.S.; N.S.M., T.S. and N.S. designed and performed the NADPH diaphorase experiments; A.L.T.M., N.S.M., N.N. and F.B. performed the protein synthesis fluorescent assay; B.H. and T.S. performed the amperometric experiments and analysed the results; A.L.T.M. and B.H. wrote the paper with comments from all the authors.

## Competing interests

The authors declare no competing interests.

## Methods

### Animals and electrophysiological recordings

Mature *Octopus vulgaris* of both sexes were collected by local fisherman on the Israeli Mediterranean coast. The conditions under which the animals were kept, the slice preparation procedures and the extracellular local field potential (LFP) recording methods followed Shomrat et al.^5^ unless mentioned otherwise. In some experiments we used isolated brain preparations that included the intact supraesophageal brain mass^4^. The fPSP of the synapses between SFL axons and AM cells was evoked by stimulating the SFL tract at 0.03 Hz. We induced LTP by high frequency stimulation (HF; 4 trains of 20 stimuli at 50 Hz, 10 s inter-train interval). Paired pulses were routinely used as test stimuli to obtain the paired pulse facilitation (PPF) ratio between the amplitudes of first and second fPSPs. The results are displayed as normalized response of the first (fPSP1/TP1) or the second (fPSP2/TP2) fPSPs. In many experiments under control conditions the first fPSP (fPSP1) was mostly indistinguishable from noise.

### NADPH diaphorase histochemistry

Immediately after removal of the supraesophageal mass from anesthetized animals the brain tissue was fixed overnight by immersion in 4% paraformaldehyde (PFA) and 0.25% glutaraldehyde in artificial seawater (ASW), pH 7.4 at 4 °C. The brain was then vibratome sectioned in the sagittal or transverse planes (50-100 µm). Tissue sections were analysed for NADPH diaphorase activity according to Hope & Vincent^51^ (modified by Moroz^29^). Briefly, slices were incubated at room temperature (RT) in the dark with reaction solution (0.5 M Tris-HCl buffer solution, pH 7.6, 1 mM β-NADPH, 0.2 mM Nitro Blue Tetrazolium (NBT), 0.3 % Triton X-100) for 30 min, rinsed with Tris-HCl buffer, then transferred to 4% PFA in methanol for one hour at RT. Finally, slices were dehydrated in ethanol, cleaned in xylene, mounted on SuperFrost Plus slides (Menzel Glaser, Germany). Specificity of NADPH diaphorase staining was tested in control experiments, in which tissue sections were incubated in the reaction solution as described, except that β-NADPH or NBT was omitted.

### FUNCAT detection of protein synthesis

To evaluate anisomycin suppression of *de novo* protein synthesis in the octopus brain, we adapted the method of fluorescent noncanonical amino acid tagging (FUNCAT) using L-azidohomoalanine (Thermo Fisher Scientific) described in detail for hippocampal slices^52^. We adapted the method aiming to guarantee marine invertebrate physiological conditions prior to fixation (ASW instead of Ringer solution, pH, and incubation temperature). Brain slices were obtained as for electrophysiological recordings. Alexa Fluor 488 azide (Thermo Fisher Scientific) was used as fluorophore-alkyne-tag. Control slices were incubated with L-azidohomoalanine in the absence of anisomycin. Confocal imaging of the treated slices was performed using a Nikon C1 confocal system mounted on a Nikon TE-2000 Eclipse microscope system with a Nikon plan-Apo 60X NA 1.4 objective. Images were collected and processed using EZ-C1 software (Nikon). Alexa Fluor 488 azide was excited with the 488 nm laser and the emission was collected with a 515 ± 30 nm filter. Images were prepared using NIH ImageJ software (Bethesda, MD, USA).

### NO amperometry

NO was measured directly in the VL using electrochemical methods with 30 µm polypropylene-insulated carbon fibre microelectrodes (CFMs) (ProCFE Dagan Corporation) coated by dipping the electrode tip five times into Nafion® 117 solution and drying the electrode for 10 min at 85 °C^34–35^. This procedure was repeated ten times. The CFMs were then coated by electropolymerization with o-phenylenediamine dihydrochloride^35^ and cellulose^53^. All these coatings were found to be essential for increasing the sensitivity and specificity of the CFM to NO in ASW. To test the electrodes, we measured the release of NO from a known concentration of NO donor (stock solution of SNAP prepared in 100 µM EDTA). 10 µM CuCl2 was added to the ASW to catalyse SNAP decomposition to NO and disulfide by-product^54^. This concentration of Cu^2+^ had no significant effect on the physiological responses. For testing the peak oxidation voltage of NO in ASW, a slowly rising (∼2.3 s) voltage ramp from a DC holding voltage of 0.5 V to 1.1 V was used as a command for clamping the CFM potential with a Chem-Clamp Voltmeter-Amperometer (Dagan Corporation). Adding 10 µM SNAP to ASW containing 10 µM CuCl2 caused a maximal positive current response close to 750 mV. Hence, for the *in vitro* measurements of NO during physiological experiments the electrode potential was clamped at a constant 750 mV. Because the oxidation current declined gradually in the presence of constant NO concentration, using a micrometric manipulator, we inserted the CFMs into the tissue (∼100 µm deep) every 5 min for only 90 s. Amperometric measurements were stored together with the recording of the LFP generated by stimulation of the SFL tract. For averaging population results (Fig. 2h) each experiment was normalized to the maximal response and aligned with respect to the last current measurement following electrode withdrawal.

### Drugs

Drugs were administered via the perfusion system, taking ∼1min to reach the recording site. We used L-NNA (*N*ω-Nitro-L-arginine, 10 mM), L-NAME (*N*ω-Nitro-L-arginine methyl ester hydrochloride, 10 mM), D-NAME (*N*ω-Nitro-D-arginine methyl ester hydrochloride, 10 mM), ODQ (1H-[1,2,4] Oxadiazolo[4,3-a]quinoxalin-1-one; 50μM), methylene blue (0.1 mM), dBcGMP (0.2mM), 8-Br-cGMP (0.1mM), 8-pCPT-cGMP (10-100μM), ZAP (Zaprinast; 0.1mM), IBMX (3-Isobutyl-1-methylxanthine; 0.5-1mM), SNAP (S-Nitroso-N-acetyl-DL-penicillamine; 1-100 µM), DETA (Diethylenetriamine NONOate; 0.4 mM), DEA (Diethylamine NONOate; 0.1mM), and PTIO (2-Phenyl-4,4,5,5 tetramethylimidazoline-1-oxyl 3-oxide; 1 mM). Some drugs were dissolved in ASW and were freshly made before the experiments (L-NNA, L-NAME, D-NAME, methylene blue). Stock solution aliquots were prepared in DDW (dBcGMP), in DMSO 0.1-0.25% (ODQ, ZAP, IBMX), in ethanol 1% (PTIO), or in NaOH 0.05 mM (DEA), and were diluted in ASW before the experiments to a concentration in which the vehicles had no physiological effects. When required, pH was adjusted to 7.6 using NaOH or HCl. The effects of pharmacological manipulations were always compared to interleaved control experiments using the drug vehicle at the same concentration. Slices treated with reversible drugs, which could be washed out, could be used a second time at a different recording location to guarantee a different and unstimulated local population of cells. All drugs and chemicals were purchased from Sigma-Aldrich, unless stated otherwise.

### Data analysis

Electrophysiological data were analysed as described in Shomrat et al.^5^. Unless stated otherwise, we used averages of 5 trials for all the electrophysiological recordings (30 s interstimulation interval). In the experiments where the slices were exposed to the drug before LTP induction, the responses were normalized relative to the control baseline (first 7.5 min of the recording). Where we sought to quantify the degree of LTP blockage, LTP was induced prior to exposure to the drug and the responses were normalized relative to LTP (7.5 min recording after HF). Values are presented as mean ± s.e.m. Paired pulse facilitation (PPF) index was calculated using to the formula (fPSP_2_/TP_2_-fPSP_1_/TP_1_) / fPSP_2_/TP_2_, which allowed quantifying PPF value also when the first fPSP amplitude was indistinguishable from the noise.

## Notes

### Competing Interest Statement

The authors have declared no competing interest.

### Summary of Updates

correcting legend labeling of the graphs in Figure 4B

